# Noise propagation in an integrated model of bacterial gene expression and growth

**DOI:** 10.1101/246165

**Authors:** Istvan T. Kleijn, Laurens H. J. Krah, Rutger Hermsen

## Abstract

In bacterial cells, gene expression, metabolism, and growth are highly interdependent and tightly coordinated. As a result, stochastic fluctuations in expression levels and instantaneous growth rate show intricate cross-correlations. These correlations are shaped by feedback loops, trade-offs and constraints acting at the cellular level; therefore a quantitative understanding requires an integrated approach. To that end, we here present a mathematical model describing a cell that contains multiple proteins that are each expressed stochastically and jointly limit the growth rate. Conversely, metabolism and growth affect protein synthesis and dilution. Thus, expression noise originating in one gene propagates to metabolism, growth, and the expression of all other genes. Nevertheless, under a small-noise approximation many statistical quantities can be calculated analytically. We identify several routes of noise propagation, illustrate their origins and scaling, and establish important connections between noise propagation and the field of metabolic control analysis. We then present a many-protein model containing > 1000 proteins parameterized by previously measured abundance data and demonstrate that the predicted cross-correlations between gene expression and growth rate are in broad agreement with published measurements.

---

Few processes are more fundamental to life than the growth and proliferation of cells. Bacterial cells in particular are highly adapted to grow rapidly and reliably in diverse habitats [1]. Yet, the composition of individual bacteria grown in a constant environment is known to fluctuate vigorously, in part due to the stochastic nature of gene expression [2–5]. Many experimental and theoretical studies have shed light on the origins, characteristics and consequences of this “noisy” expression [2–17]. Still, it remains unknown to what extent, and by what routes, noise in gene expression propagates through the cell and affects the rate of growth [5,18,19], which is often considered a proxy for its fitness [18, 20].

Recently, important progress towards understanding noise propagation in single cells has been made through experiments in which the instantaneous growth of individual *Escherichia coli* cells was monitored in real time under fixed growth conditions [5, 21]. Such experiments have revealed large fluctuations in the growth rate, with coefficients of variation of the order of 25%, which in part result from noise in the concentrations of metabolic enzymes [5]. Conversely, growth-rate fluctuations affect the concentrations of individual enzymes, because the cell’s constituents are diluted whenever the cell grows [22]. Such results emphasize that a clear understanding of these processes is complicated by the fact that gene expression, metabolism, and growth are highly interdependent, involving multiple layers of feedback and cellular constraints.

This interdependence is also central to a series of recent studies that characterize the *average* composition and growth rate of *Escherichia coli* cultures in balanced exponential growth under variation of the growth medium [23–29]. In particular, these experiments have revealed striking linear relations between their mean proteomic composition and their mean growth rate [26–31]. Phenomenological models have demonstrated how such “growth laws” can be understood as near-optimal solutions to constrained allocation problems [20, 32–34]. These results also stress that global physiological variables and constraints strongly affect the expression of individual genes. As such, both these experiments and the single-cell experiments mentioned above suggest a “holistic” perspective: the behavior of individual components cannot be understood without some knowledge of the cell’s global physiological state [35, 36].

Here, we present a model of bacterial cells growing under fixed external growth conditions, in which gene expression, metabolism and growth are fully integrated. We offer a highly simplified description that nevertheless imposes several essential global cellular constraints. Both gene expression and growth rate fluctuate due to the stochastic synthesis of many protein species that together control the rates of metabolism and growth. Conversely, the rate of metabolism constrains the protein synthesis rates and the growth rate sets the dilution rate of all proteins. As a result, noise in the expression of each gene propagates and affects the expression of every other gene as well as the growth rate—and *vice versa*.

Below, we first introduce the generic modeling framework and its assumptions. We then make an excursion to the theory of growth control, in order to define growth-control coefficients and establish connections between the propagation of noise and the field of Metabolic Control Analysis. Next, we discuss how the concentration of each protein is affected by the synthesis noise in all other proteins; this exposes a hidden assumption in a standard operational definition of intrinsic and extrinsic expression noise. We subsequently explain the noise modes that characterize the noise propagation between gene expression and growth in the context of a toy model with just two proteins. Lastly, we present a many-protein model that includes 1021 protein species with experimentally measured parameters. We demonstrate that the cross-correlations functions between expression and growth rate predicted by this model capture the main features of published measurements.

## Results

### Modeling framework

We here discuss the key assumptions of the modeling framework (Fig 1); see S1 Text, pp. 1–6 for details. We consider a culture of bacterial cells that has reached steady-state exponential growth under fixed external growth conditions. We study fluctuations of gene expression within individual cells in this steady state, and in particular how these fluctuations reverberate through the growing cell. Similar assumptions connecting the increase in biomass, the cellular growth rate, protein synthesis, and growth-mediated dilution were explored in a recent review article [37].

**Fig 1.**
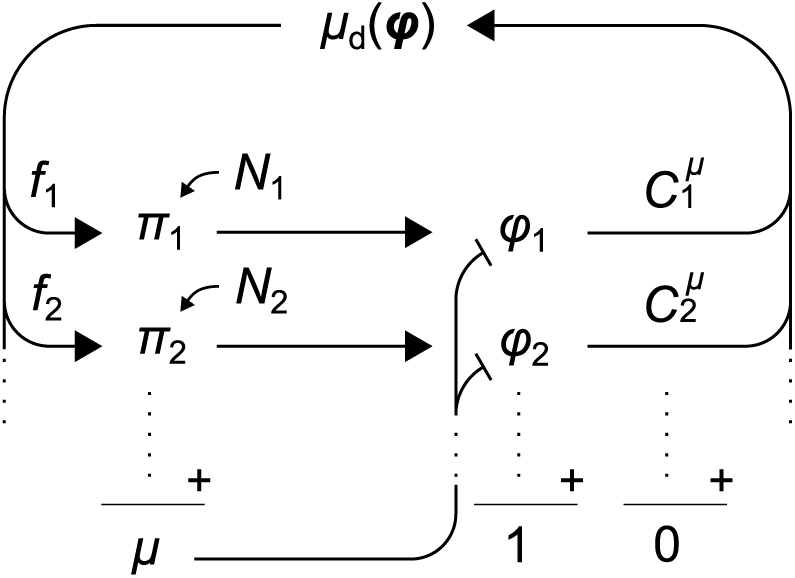
Integrated model of stochastic gene expression and cell growth. The cell contains many protein species, with proteome mass fractions *ϕ*_*i*_ that sum to 1. Mass fractions are increased by protein synthesis but diluted by growth. The synthesis rate *π*_*i*_ of each species *i* is modulated by a noise source *N*_*i*_. The instantaneous growth rate *µ* reflects the total rate of protein synthesis. Proteins affect metabolism and thus the deterministic growth rate *µ*_d_(*ϕ*), as quantified by growth-control coefficients 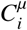. A fraction *f*_*i*_ of the total metabolic flux is allotted to the synthesis of protein *i*. The inherent noise in the expression of each gene reverberates through the cell, affecting cell growth and the expression of every other gene.

The mass density of *E. coli* cells is dominated by protein content [38] and under tight homeostatic control [39]. We assume that this homeostasis also eliminates long-lived proteindensity fluctuations in single cells. Then, the volume of a cell is proportional to its protein mass *M*: = ∑_*i*_ *n*_*i*_, where *n*_*i*_ is the abundance (copy number) of protein *i*. (We ignore that different proteins have different molecular weights.) The instantaneous growth rate is then defined by *µ:= M/M*, and the proteome fraction *ϕ*_*i*_:= *n*_*i*_/M of enzyme *i* measures its concentration. Differentiation of *ϕ*_*i*_ with respect to time then yields

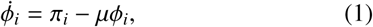

where *π*_*i*_ is the synthesis rate per protein mass. (Here we neglect active protein degradation, which on average amounts to about 2% of the dilution rate [40].) By definition, proteome fractions obey the constraint ∑_*i*_ *ϕ*_*i*_ = 1. Combined with Eq. (1) this results in

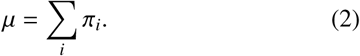

That is, the growth rate equals the total rate of protein synthesis.

Another key assumption of our model is that the cellular growth rate is an *intensive* quantity. That is: given fixed mass fractions, the growth rate does not depend on the cell size, as suggested by the observation that individual *E. coli* cells grow approximately exponentially within their cell cycle [5, 41]. Based on this, we express the synthesis rate of protein *i* as:

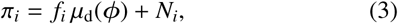

in which

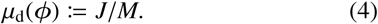

The first term in equation (3) is an intensive function; it captures the deterministic effect of the cellular composition *ϕ* = (*ϕ*_1_*, ϕ*_2_, …) on the metabolic flux *J* that quantifies the rate of biomass production, normalized by the protein mass *M*. (Note that, here and below, we use the term metabolism in a broad sense; it is intended to encompass all catabolic and anabolic processes required for biomass production and cell growth, including protein synthesis.) The coefficients *f*_*i*_ specify which fraction of this flux is allocated towards the synthesis of protein species *i*. Because the *f*_*i*_ are fractions, ∑_*i*_ *f*_*i*_ = 1.

The second term of equation (3) couples each synthesis rate *π*_*i*_ to a zero-mean Ornstein–Uhlenbeck noise source *N*_*i*_ that represents the stochasticity of both transcription and translation [42]. Each noise source is characterized by an amplitude *θ*_*i*_ and a rate of reversion to the mean *β*_*i*_; the latter’s inverse 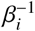 characterizes the time scale of intrinsic fluctuations in *π*_*i*_. The variance of *N*_*i*_ is given by Var 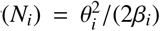. All noise sources are mutually independent, and we neglect other sources of noise, such as the unequal distribution of molecules over daughter cells during cell division (see Discussion).

Combining equation (2) and (3) reveals that

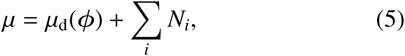

which identifies *µ*_d_(*ϕ*) as the growth rate afforded by a given proteome composition *ϕ* in the zero-noise limit. Given a function *µ*_d_(*ϕ*), equations (1)–(3) fully define the dynamics of the cell.

Below, we focus on the simplest case where, under given environmental conditions, the allocation coefficients *f*_*i*_ are constant. This means that the cell does not dynamically adjusts its allocation in response to fluctuations in expression levels. We note, however, that such dynamical effects of gene regulation could be included by allowing the *f*_*i*_ to depend on intraand extra-cellular conditions, and in particular on the cellular composition *ϕ*. (See S1 Text, p. 4.) We also stress that the allocation coefficients may differ strongly between growth conditions, as demonstrated by the growth laws mentioned above. For example, the *f*_*i*_’s of ribosomal proteins must be considerably larger in media that support a fast growth rate than in media with strong nutrient limitation, because the mean mass fraction of ribosomal proteins increases with the growth rate [30]. Here, however, we describe stochastic cell growth under fixed environmental conditions, so that the (mean) allocation of resources is well-defined and knowable in principle—for example through proteomics data.

Fig 1 is an illustration of the modeling framework. Noise in the synthesis of a protein species induces fluctuations in its mass fraction (equation (1)). Through their effect on metabolism, these fluctuations propagate to the deterministic growth rate *µ*_d_, which modulates the synthesis of all protein species (equation (3)). In parallel, all noise sources directly impact the growth rate *µ* (equation (5)) and thus the dilution of all proteins (equation (1)).

### Linearization under a small-noise approximation

The results below rely on the assumption that equations (1)–(5) may be linearized around the time-averaged composition *ϕ***_0_**. This transforms equation (5) to

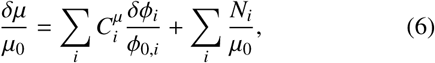

where *δϕ*_*i*_ is the deviation of *ϕ*_*i*_ from its time average *ϕ*_0*,i*_ and *δµ* the deviation of *µ* from *µ*_0_ *µ*_d_(*ϕ***_0_**). (See Text S1, p. 3 for derivations.) The coefficients *C*^*µ*^ are defined as

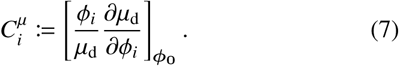

In the terminology of linear noise models, the 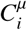 are transfer coefficients: they quantify to what extent fluctuations in *ϕ*_*i*_ transmit to *µ*_d_. Equation (6) demonstrates that the growth rate is affected by all noise sources, both directly (second term on the right-hand side) and indirectly through fluctuations in the protein mass fractions.

### Transfer coefficients are growth-control coefficients

The transfer coefficients 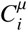 are reminiscent of the logarithmic gains defined in biochemical systems theory, which relate enzyme abundances to the metabolic flux in a given pathway [43]. It has previously been shown that these gains are relevant in the context of noise propagation [44]. Here, however, we consider the growth rate of the cell rather than the flux through a distinct pathway. In this section, we connect the transfer coefficients 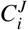 to the control of cellular growth and the field of Metabolic Control Analysis (MCA) [45, 46].

In MCA, flux-control coefficients (FCCs) 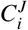 are defined that quantify to what extent an enzyme concentration *ϕ*_*i*_ limits (controls) a metabolic flux *J i* [45, 46]:

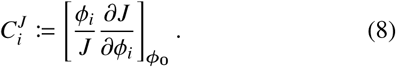

In direct analogy to this definition of FCCs, the transfer coefficients of equation (7) can be interpreted as growth-control coefficients (GCCs) that quantify each enzyme’s control of the growth rate. From equation (4) a direct link between FCCs and GCCs can be derived (see also [47], p. 7 of S1 Text, and S1 Fig):

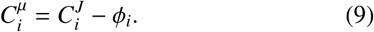

The GCCs are specified by the sensitivity of the growth rate *µ*_d_(*ϕ*) to changes in the proteome composition *ϕ*, evaluated in the steady-state mean, *ϕ***_0_**. Both the mean composition *ϕ***_0_** and the function *µ*_d_ clearly differ between growth conditions; therefore, the GCCs depend on the growth conditions as well.

As mentioned, studies on the resource allocation of cells grown under different growth conditions have revealed striking empirical relations between the mean proteome composition and the mean cellular growth rate [26, 28–30]. Even though these growth laws describe relations between growth rate and composition, they should not be confused with *µ*_d_. The growth laws describe correlations between the mean composition and the mean growth rate under variation of the growth conditions, whereas *µ*_d_ describes the deterministic effect of the instantaneous composition on the instantaneous growth rate under a particular, fixed growth condition. There is no direct relation between the two. By extension, the growth laws do not directly translate into knowledge on the GCCs.

### Growth-control coefficients and their sum rule

An important difference between metabolic flux and cellular growth rate lies in their behavior under a scaling of the system size. It is routinely assumed that metabolic fluxes scale linearly with the system size, meaning that an increase in the abundances of all enzymes by a factor *α* increases the metabolic flux *J* by the same factor *α*. That is, fluxes are *extensive* variables. Based on this assumption, a famous sum rule has been derived for FCCs [45, 46]:

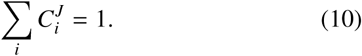

In contrast, we assumed the growth rate to be invariant under scaling of the system size, *i.e*, that the growth rate is an *intensive* variable. (Indeed, as equation (4) directly shows, if *J* is extensive, *µ*_d_ must be intensive, and *vice versa*.) Under this assumption, GCCs obey a markedly different sum rule:

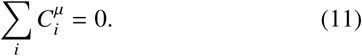

This sum rules articulate a delicate trade-off: the excess of one protein implies the lack of another.

Both sum rules are special cases of Euler’s homogeneous function theorem. Specific derivations are presented in S1 Text on p. 7. In general, for an arbitrary function *f* with a scaling relation *f* (*αϕ*) = *α*^*k*^ f (*ϕ*), a sum rule can be derived by differentiating this equation with respect to *α* and evaluating the result in *α* = 1. The particular cases *k* = 1 (for the flux *J*), and *k* = 0 (for the growth rate *µ*_d_) lead to equations (10) and (11).

In theory, all expression levels could be regulated such that 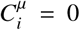 for all protein species *i*. In reality, however, many protein species do not have a function within metabolism or biomass growth. By definition, the metabolic flux *J* does not depend on the expression levels of these proteins; therefore, their FCCs are zero. The GCC of such a protein, with mass fraction *ϕ*_*h*_, then follows from equation (9):

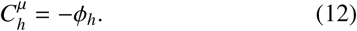

That is, the control of all non-metabolic enzymes on the growth rate is negative. The sum rule then implies that the sum of GCCs of all proteins that do contribute to biomass growth must be positive and equal to

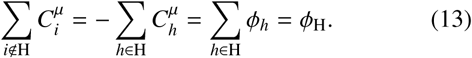

where H denotes the set of non-metabolic proteins. This goes to show that any system that bears the cost of producing nonmetabolic proteins must contain other proteins that have positive growth control.

This conclusion has implications for the propagation of noise. We saw that the the noise transfer coefficients appearing in the linear noise model are in fact GCCs. The analysis in the previous paragraph demonstrates that these GCCs cannot all vanish; it then follows that there must be linear-order noise transfer from protein levels to the growth rate in all cells that maintain non-metabolic proteins.

Non-metabolic proteins are common, both in wild-type cells and in engineered constructs. In wild-type *E. coli*, the expression level of proteins that do not contribute to biomass growth were estimated recently in a study that combined a genome-scale allocation model with proteomics data sets [48]. Direct estimates of *ϕ*_H_ ranged from 25% to 40%, depending on the precise growth conditions. Although not directly beneficial to the growth of the cell in constant environments, the non-contributing proteome fraction is thought to provide fitness benefits to cells that encounter frequent changes in growth conditions [48]. Furthermore, synthetic biologists commonly study systems with a large expression burden [49].

### Separating in- and extrinsic noise components

Within the above framework, many statistical properties can be calculated analytically [5, 42]. In particular, the noise level of the concentration of protein *i*, quantified by the coefficient of variation?_*i*_, can be expressed as:

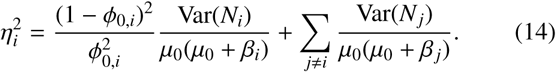

The derivation is provided in S1 Text, pp. 4–6. Equation (14) shows that the coefficient of variation has two components: the first term results from the noise in the synthesis of the protein itself, the second from the noise in the synthesis of all other proteins. Each term is proportional to the variance of the corresponding noise source, but weighted by a factor that decreases with the mean growth rate *µ*_0_ and the reversion rate *β*_*i*_ of that noise source. This analysis confirms that the inherent noise in the synthesis of one protein affects all other proteins.

A fundamental distinction is commonly made between intrinsic and extrinsic noise in gene expression [44]. Intrinsic noise results from the inherently stochastic behavior of the molecular machinery involved in gene expression; extrinsic noise from fluctuations in the intra- and extracellular environment of this machinery. In this sense, the two terms in equation (14) can be identified as intrinsic and extrinsic contributions.

Complications arise, however, if the standard operational definition of these terms is applied [4, 6]. This definition considers two identical reporter constructs R and G expressed in the same cell (Fig 2A). Noise sources extrinsic to both reporters affect both reporters identically, inducing positively correlated fluctuations in the concentrations of the reporter proteins. Intrinsic noise sources instead produce independent fluctuations in each concentration. Extrinsic noise is therefore measured by the covariance between both expression levels; intrinsic noise by their expected squared difference. This operationalization, however, implicitly assumes that intrinsic noise does not propagate between the reporters. This assumption is violated in our model because the synthesis of reporter R directly contributes to the dilution of protein G (Fig 2B). Consequently, the covariance between the expression levels has two contributions:

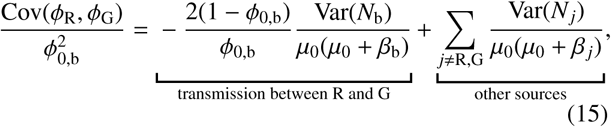

where the label “b” indicates quantities that are by definition identical for both expression systems. The second term on the right-hand side is positive and stems from noise sources that affect both reporters identically. The first term, however, is negative; it reflects the transmission of noise between reporters R and G. It would be misleading to identify equation (15) as the extrinsic component of the noise—it is not even guaranteed to be positive. We conclude that the operational definition is not suitable when noise propagates between arbitrary genes.

**Fig 2.**
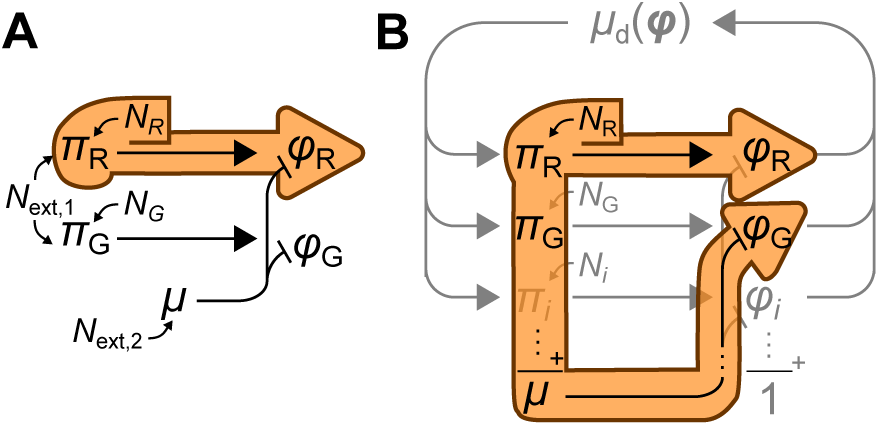
Limitations of the operational definition of inand extrinsic expression noise. (*A*) Extrinsic noise is measured by the covariance between the expression levels of two identical reporter systems R and G. This presupposes that the intrinsic noise *N*_R_ of system R affects concentration *ϕ*_R_ but not *ϕ*_G_ (orange outline), so that the covariance between *ϕ*_R_ and *ϕ*_G_ quantifies the contribution of extrinsic sources *N*_ext*,i*_. (*B*) But in our model, *N*_R_ affects the growth rate and thus the dilution of *ϕ*_G_. This adds a negative term to the covariance, which no longer measures just the extrinsic noise.

### Expression–growth correlations in a two-protein toy model

The circulation of noise in the cell can be studied by measuring cross-correlations between expression and growth rate in single-cell experiments [5]. Interpreting measured crosscorrelations, however, is non-trivial. To dissect them, we now discuss a toy version of the model with just two protein species, X and Y. Despite its simplicity, it displays many features seen in more realistic models.

Within the linear noise framework, *ϕ*_Y_–*µ* and *π*_Y_–*µ* crosscorrelations, respectively denoted *R*_*ϕ*Y *µ*_(*τ*) and *R*_*π*Y *µ*_(*τ*), can be calculated analytically [42]. Up to a normalization, the results can be written as:

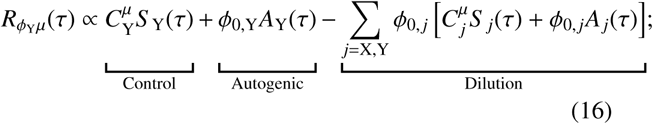

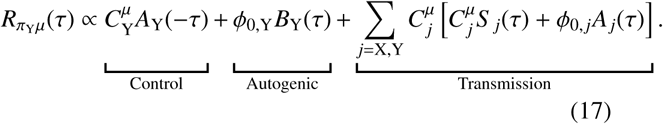

(For a full derivation, not limited to the two-protein case, see S1 Text, pp. 5–6. The two-protein case is discussed further in S1 Text, pp. 8–9.) These equations are plotted in Fig 3AB (see caption for parameters). As the equations show, the crosscorrelation functions are linear combinations of three functions *S*_*i*_(*τ*), *A*_*i*_(*τ*), and *B*_*i*_(*τ*), which are also illustrated in the figure.

**Fig 3.**
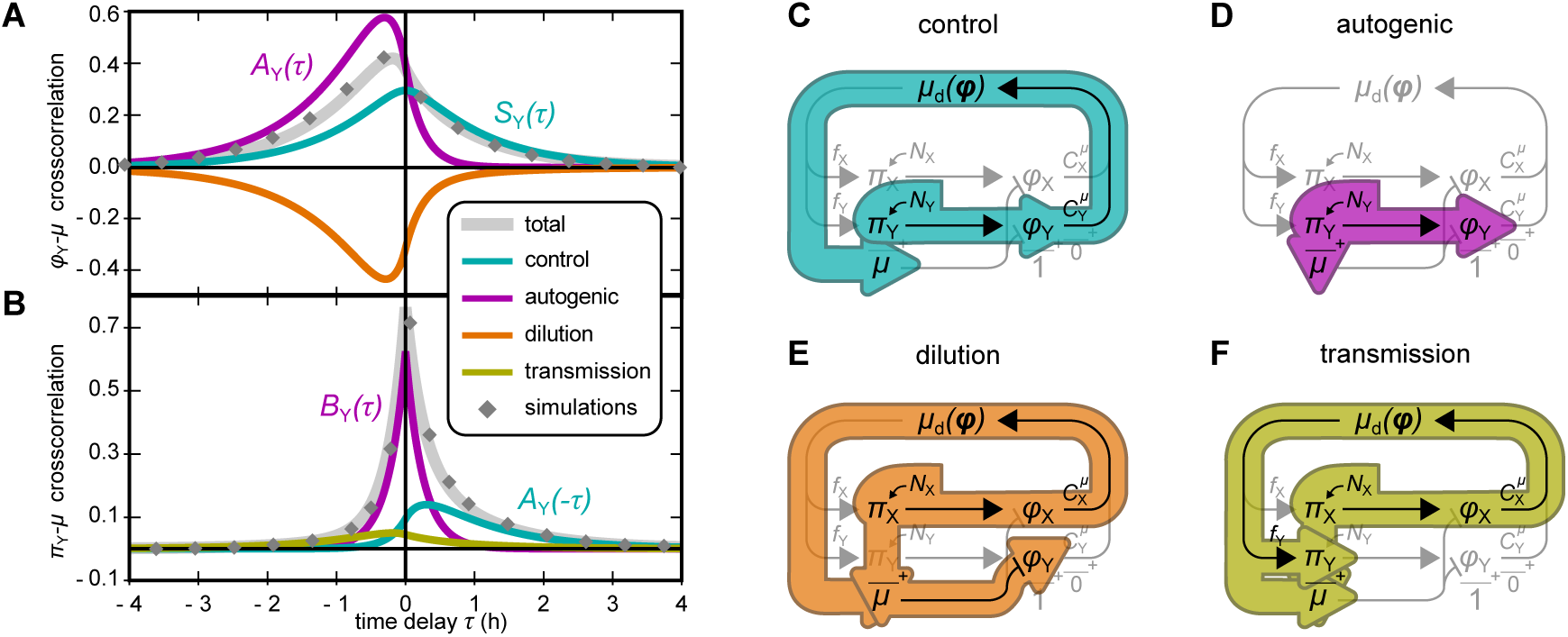
Noise modes in a toy model containing only two protein species, X and Y. (*A*) Analytical solution for the cross-correlation between protein Y’s proteome fraction *ϕ*_Y_ and growth rate *µ* (gray curve), verified by simulations (gray diamonds, details in S1 Text, p. 9). The contributing noise modes are indicated (colored curves). (*B*) Same as (A), but for the synthesis rate *π*_Y_. The cross-correlation functions are linear combinations of three classes of functions, called *A*_*i*_(*τ*), *B*_*i*_(*τ*), and *S* _*i*_(*τ*) (see S1 Text, equations (47)–(49) for their definitions). In panels (A) and (B), noise modes that are proportional to just one of these functions are annotated accordingly. (*C*)–(*F*) Noise propagation routes underlying the noise modes. The control mode and the autogenic mode arise from noise source *N*_Y_ alone. Both noise sources *N*_X_ and *N*_Y_ contribute to the dilution and transmission modes, but only the contribution of *N*_X_ is illustrated in Fig (D) and (F). Parameters for (A) and (B): 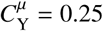; *ϕ*_0,Y_ = 0.33; mean growth rate *µ*_0_ = 1 h^−1^; noise sources of *N*_Y_ and *N*_X_ have amplitudes *θ*_Y_ = 0.5 and *θ*_X_ = 0.5 and reversion rates *β*_Y_ = *β*_X_ = 4*µ*_0_.

To aid interpretation, the cross-correlations can be decomposed into four noise modes, as indicated in equations (16) and (17).

The **control mode** (Fig 3C) reflects the control of enzyme Y on the growth rate. Noise *N*_Y_ in the synthesis of Y causes fluctuations in *ϕ*_Y_, which transfer to the growth rate in proportion with the 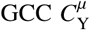. Because the effect of *ϕ*_Y_ on *µ* is instantaneous, the contribution to the *ϕ*_Y_–*µ* cross-correlation is proportional to the *symmetric* function *S* _Y_(*τ*). In contrast, the effect of *π*_Y_ on *µ* involves a delay; hence the contribution to the *π*_Y_–*µ* cross-correlation is proportional to the *asymmetric* function *A*_Y_(*τ*). In both cases, the amplitude scales with 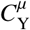.

The **autogenic mode** (Fig 3D) is a consequence of equation (2). Because the growth rate matches the total rate of protein synthesis, noise in the synthesis of Y instantly affects the growth rate, resulting in a noise mode in the *π*_Y_–*µ* cross-correlation that is proportional to the symmetric function *B*_Y_(*τ*). With a delay, this noise also affects *ϕ*_Y_, adding an asymmetric mode to the *ϕ*_Y_–*µ* cross-correlation. This mode does not depend on the control of Y; instead, its amplitude is proportional to the mean concentration *ϕ*_0,Y_.

The **dilution mode** (Fig 3E) pertains only to the *ϕ*_Y_–*µ* cross-correlation. It reflects that the growth rate of the cell is also the dilution rate of protein Y (equation (1)). With a delay, upward fluctuations in *µ* therefore cause downward fluctuations in *ϕ*_Y_. A subtle complication is that noise in the synthesis rate of both proteins reaches *µ* via two routes: through the immediate effect of *π*_Y_ on *µ*, and through the delayed effect of *π*_Y_ on *ϕ*_Y_, which in turn affects *µ* in proportion with 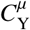 (see in equation (6)). Together, these routes result in a mode towards which each protein contributes both a symmetric and an asymmetric function.

Lastly, the **transmission mode** (Fig 3F) is unique to the *π*_Y_–*µ* cross-correlation. It reflects that all noise sources affect the cell’s composition *ϕ* and therefore *µ*_d_; this in turn induces fluctuations in the synthesis rate *π*_Y_. The noise sources again affect the growth rate via the two routes explained above, causing a symmetric and an asymmetric component to the *π*_Y_–*µ* cross-correlation for each protein.

The above analysis shows that, even in a highly simplified linear model, the cross-correlations are superpositions of several non-trivial contributions. The intuitions gained from this exercise will be used below when we present the results of a more complex model.

### The effects of gene regulation

Above, we assumed that the cell allocates a fixed fraction *f*_*i*_ of its metabolic flux towards the synthesis of protein *i*. Within this two-protein model all cross-correlations can still be computed if the *f*_*i*_ are linear(ized) functions of the concentrations *ϕ* (see S1 Text, pp. 8–9, and S2 Fig). The resulting feedback regulation affects the decay of fluctuations: a negative feedback shortens the correlation time scales and reduces variance, whereas positive feedback lengthens them and increases variance (*cf.* [3, 10, 50]).

### Expression–growth correlations in a many-protein model

In single *E. coli* cells, the cross-correlations between gene expression and growth rate have been measured by Kiviet *et al.* [5]. To test whether the above framework can reproduce their results, we constructed a model that includes 1021 protein species with realistic parameters, based on an experimental data set [51].

In the experiments, micro-colonies of cells were grown on lactulose (a chemical analog of lactose) and expression of the *lac* operon was monitored using a green fluorescent protein (GFP) reporter inserted in the operon. Because intrinsic fluctuations in GFP expression affect the cross-correlations directly as well as indirectly, through their impact on the growth rate and the expression of other genes, we modeled this reporter construct explicitly (see Fig 4A, and S1 Text, pp. 9–11). Specifically, the *lac* operon O was represented as a collection of three proteins Y, Z, and G (for LacY, LacZ, and GFP) affected by a shared noise source *N*_O_ in addition to their private sources *N*_Y_, *N*_Z_, and *N*_G_. The GCC of the operon as a whole is the sum of the GCCs of its genes.

**Fig 4.**
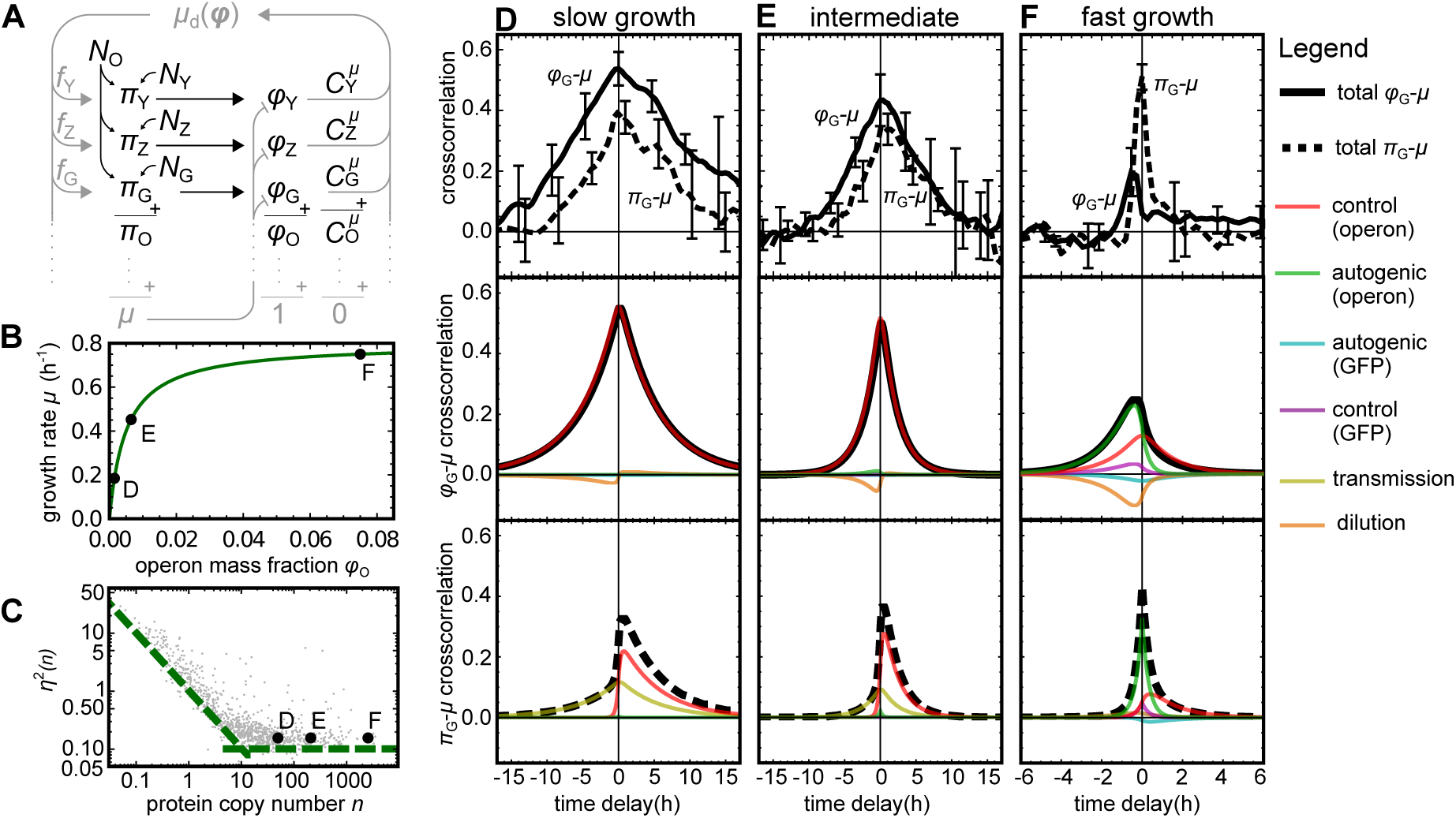
Expression–growth cross-correlations in the many-protein model. (*A*) Cartoon of the noise propagation network. (*B*) Monod curve describing the mean growth rate as a function of *lac* expression. Black dots indicate the operon mass fractions and growth rate used to calculate the cross-correlations in (D)-(F). (*C*) Noise distribution of the proteome (gray cloud) taken from Ref. [51], and the values chosen for proteins on the *lac* operon (black dots). Green dashed lines are guides for the eye. (*D*)–(*F*) Experimental [5] (top panels) and theoretical (middle and bottom panels) cross-correlations for three growth conditions. Proteome fraction–growth and production–growth cross-correlations are plotted as solid and dashed black lines, respectively. As in Fig 3AB, colored lines show the contributing noise modes.

By varying the mean expression of the *lac* operon with a synthetic inducer, Kiviet *et al* measured cross-correlations in three growth states with different macroscopic growth rates: “slow”, “intermediate”, and “fast” [5]. Empirically, the macroscopic growth rate obeyed a Monod law [52] as a function of the mean *lac* expression. We therefore mimicked the three growth states by choosing their mean *lac* expression levels and growth rates according to three points on a Monod curve that approximates the empirical one (Fig 4B, labels D, E, and F). Via equation (7), the same curve also is also used to estimate the GCC of the *lac* operon in each condition. Under “slow” growth conditions, the *lac* enzymes limit growth considerably (large GCC); under “fast” conditions, *lac* activity is almost saturated (small GCC).

To choose realistic parameter values for all other proteins, we used a published dataset of measured means and variances of *E. coli* protein abundances [51]. For each of the 1018 proteins in the dataset, the model included a protein with the exact same mean and variance (see Fig 4C). This uniquely fixed the amplitudes of all noise sources. The GCCs of all proteins were randomly sampled from a probability distribution that obeyed the sum rule of equation (11). (See Materials and Methods, and S1 Text, p. 10–11)

### Comparison with measured cross-correlations

The experimental results on the cross-correlations between GFP synthesis *π*_G_, GFP expression *ϕ*_G_, and growth rate *µ* [5] are reproduced in Fig 4D–F (top panels), together with the model predictions (middle and bottom panels).

The predicted cross-correlations are linear superpositions of the same noise modes as described for the two-protein model. However, the dilution and transmission modes are now driven by all 1022 noise sources, and there are two instances of the control and autogenic modes: one associated with the expression and GCC of the operon as a whole, and one with the expression and GCC of GFP separately. (See equations (89)–(94) in S1 Text, p. 10.)

At slow growth, the *ϕ*_G_-*µ* cross-correlation is almost symmetrical (Fig 4D, middle panel). Here the control mode of the operon dominates due to its large GCC. At higher growth rates, the autogenic modes become more prominent because their amplitudes are proportional to the expression level of the *lac* genes; at the same time, the amplitudes of the control modes decrease with the GCCs (Fig 4EF, middle panels). As a result, the cross-correlation becomes weaker and more positively skewed.

At slow growth, the *π*_G_–*µ* cross-correlation is negatively skewed because the operon control mode is dominant (Fig 4D, bottom panel). It also shows a notable transmission mode. With increasing growth rate, the autogenic modes increase in importance, which narrows the peak, increases its height, and reduces its asymmetry (Fig 4EF, bottom panels). The patterns seen in both cross-correlations are in good qualitative agreement with the experimental data (Fig 4DEF, top panels).

### Alternative dataset, similar results

In the dataset that we used to parameterize protein expression, the abundances are consistently low compared with other studies [29, 53]. However, an alternative analysis based on different abundance data [53] and sampled variances [16] yielded similar results (S1 Text p. 11, and S3 Fig). We conclude that the qualitative trends are insensitive to the precise dataset used.

## Discussion

We have presented a model of stochastic cell growth in which the growth rate and the expression of all genes mutually affect each other. Systems in which all variables communicate to create interlocked feedback loops are generally hard to analyze. Analytical results were obtained by virtue of stark simplifying assumptions. Nevertheless, the predicted and measured cross-correlations have similar shapes and show similar trends under variation of the growth rate.

That said, a few differences are observed. Chiefly, at slow and intermediate growth rates the model consistently underestimates the decorrelation timescales (peak widths). In the model, the longest timescale is the doubling time; this timescale is exceeded in the experimental data. This suggests a positive feedback that is not included in the model, possibly as a result of gene regulation (also see S2 Fig), or else a noise source with a very long auto-correlation time.

Alongside their measurements, Kiviet *et al.* published their own linear noise model, which fits their data well. In fact, the shapes of the noise modes emerging in that model are mathematically identical to those presented above [42]. Yet, the models differ strongly in their setup and interpretation. Kiviet *et al.* model a single enzyme E that is produced and diluted by growth. It features only three noise sources: one directly affects the production of E (“production noise”), one the growth rate *µ* (“growth noise”), and one affects both simultaneously (“common noise”). While these ingredients are sufficient to fit the data, the interpretation and molecular origins of the common and growth noise are left unspecified. In our model, which includes many proteins, similar noise modes emerge without explicit growth or common noise sources. Each enzyme perceives fluctuations in the expression of *all* genes as noise in the growth rate; this results in a dilution mode similar to that of Kiviet *et al*. Furthermore, noise in the synthesis of each enzyme instantaneously affects the growth rate (equation (2)) due to the assumed homeostatic control of protein density. Hence, this noise behaves as a common noise source, which explains why the autogenic mode is mathematically identical to the common-noise mode of Kiviet *et al*. We conclude that noise in the expression of many enzymes, combined with homeostatic control of protein density, can contribute to the observed but unexplained commonand growthnoise modes.

Control coefficients are routinely used in metabolic control analysis [45, 46, 54, 55] and have also been studied in the context of evolutionary optimization [47, 56]. In our linearized model, GCCs emerged as transfer coefficients, indicating that these quantities also affect the propagation of noise. Conversely, this suggests that GCCs could be inferred from noise-propagation measurements. For example, the Pearson correlation coefficient (cross-correlation at zero delay) between *ϕ*_*i*_ and *µ* might be used as an indication of control. However, we have seen in Fig 3 that the *ϕ*–*µ* correlation involves several noise modes that are independent of the GCC. As a result, the signs of the Pearson correlation and the GCC do not necessarily agree (see Fig 5A). In addition, the intrinsic noise and GCC of the reporter protein can result in a negative crosscorrelation even if the operon’s control is positive (Fig 5B). Alternatively, the asymmetry of the control mode in the *π*–*µ* cross-correlation could perhaps be exploited [5] (S4 Fig). Unfortunately, this asymmetry is also affected by other modes, such as the transmission mode, which can mask the effect (S4 Fig, panel C). We conclude that, in any case, such results have to be interpreted with great caution, ideally guided by a quantitative model.

**Fig 5.**
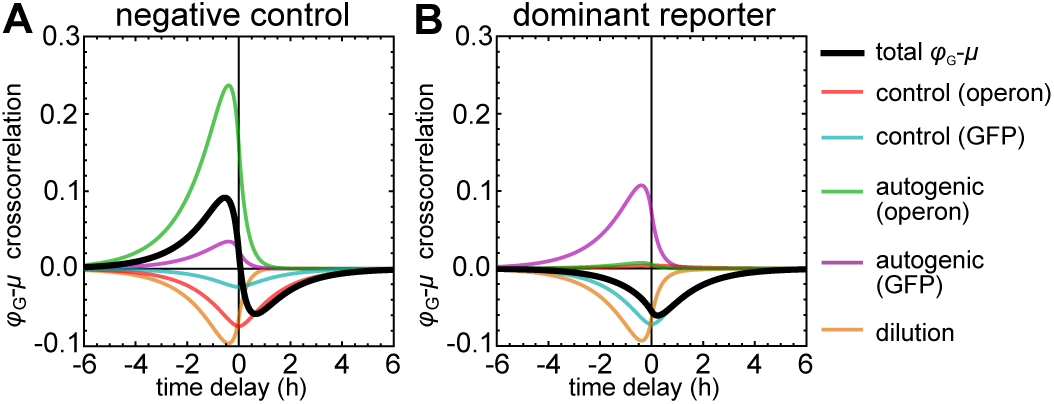
Deceptive concentration–growth cross-correlations. *(A)* Positive Pearson correlation despite a negative operon GCC, due to a dominant autogenic mode. Same parameters as Fig 4F, but with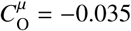. *(B)* Negative Pearson correlation despite a positive operon GCC, due to noisy GFP expression. Same parameters as Fig 4F, but with operon noise much smaller than GFP noise (see Materials and Methods).

Future theoretical work should aim to relax assumptions and remove limitations. The assumed strict control of protein density can be relaxed by allowing density fluctuations. If these are long-lived, they will likely weaken the autogenic mode and introduce new modes of their own. Also, additional noise sources can be included that do not stem directly from protein synthesis. In particular, we ignored noise originating from cell division despite its importance [8, 57, 58]. In addition, gene regulation will affect some noise modes; this can be studied by allowing the *f*_*i*_ to depend on *ϕ*. It will also be interesting to include non-protein components of the cell, such as RNAs.

A further caveat is that the linear approximation used here is only reasonable if the noise is sufficiently weak. In fact, in the presence of strong non-linearities, the approach may even break down completely. For instance, it has been shown that cellular growth can be stochastically arrested when an enzyme whose product is toxic to the cell is expressed close to a threshold beyond which toxic metabolites build up to lethal doses [59]. In such circumstances, expression level noise in those enzymes can have a highly nonlinear effect on the cellular growth rate, resulting in subpopulations of growth-arrested cells [59]. That said, under more ordinary conditions linear models that describe noise in cellular networks have previously been used to great success [5, 42].

Throughout this document we have considered noise sources that act on each production rate independently. Alternatively, one could hypothesize that the observed fluctuations in protein concentrations might instead originate from noise in the allocation of the flux—that is, from fluctuations in the allocation coefficients *f*_*i*_. This would be expected under the supposition that ribosomes are always fully occupied and translating at a constant, maximal rate, so that the relative rates of protein synthesis are determined solely by competition between different mRNAs based on their relative abundances and their translation initiation rates. Protein synthesis rates then become intrinsically correlated: an increase in the synthesis rate of one protein requires an simultaneous decrease in the synthesis rates of other proteins. In future work, such alternative models could be explored in detail. Preliminary simulations, however, show a striking symmetry in the *ϕ*_*i*_–*µ* cross-correlation and a consistent asymmetry in the *π*_*i*_–*µ* cross-correlation (for details see S1 Text pp. 12–13, and S5 Fig). This can be understood as follows. If an increase in a particular synthesis rate is always compensated by a decrease in other production rates, the noise does not affect the sum of all production rates nor the growth rate instantaneously. Therefore, no autogenic mode should be present. Notably, in our model it is the autogenic mode that explains the asymmetry in the measured *ϕ*_*i*_–*µ* cross-correlations as well as the dominant symmetric mode in the π_*i*_–*µ* cross-correlations under the fast growth condition. We conclude that noise on flux allocation alone cannot readily explain these experimental findings and additional noise sources would have to be included, such as the common noise as defined by Kiviet *et al.* [5].

Lastly, we hope that this work will inspire new experiments that can confirm or falsify the assumptions and results presented above. In particular, single-cell measurements of mass-density of protein-density fluctuations [60, 61] could establish whether our assumption of density homeostasis is warranted. Also, additional single-cell measurements could determine whether expression noise indeed propagates between reporter proteins, adding to their covariance, and whether the amplitude of the various noise modes scales with the GCCs and mass fractions as predicted.

## Materials and Methods

We here specify the parameters used for the many-protein model; also see S1 Text, pp. 10–11.

### Growth rates and protein abundances

The Monod curve (Fig 4B) is given by *µ*_0_ = *µ*_max_*ϕ*_0,O_/(*ϕ*_half_ + *ϕ*_0,O_), with *µ*_0_ the mean growth rate, *ϕ*_0,O_ the mass fraction of the *lac*-operon proteins, *µ*_max_ = 0.8 h^−1^, and *ϕ*_half_ = 0.005. The three growth states correspond to three points on this curve, with ϕ_0,O_*/ϕ*_half_ ={0.3, 1.3, 15}; this mass is shared equally among proteins Y, Z, and G. The mass fractions of the remaining proteins matched the proportions of the dataset [51].

### Ornstein–Uhlenbeck noise sources

The amplitudes of all noise sources were uniquely fixed by the constraints that (i) the CV of each Lac protein was 0.15, the amplitude of *N*_O_ was 1.5 times that of *N*_G_ [4], and (iii) all other CVs agreed with the dataset [51]. All noise reversion rates were set to *β* = 4*µ*_max_.

### GCCs

To select the GCCs, we first randomly assigned proteins (≈25% of the total mass) to the non-metabolic sector H. After the *lac* reporter construct was added, the GCC of each protein *h* ∈ H was set by equation (12). In each growth state, the GCC of the *lac* operon was calculated from the Monod curve, which yielded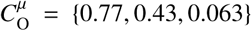. Assuming GFP is non-metabolic and the GCCs of Y and Z are equal, we set 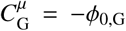 and 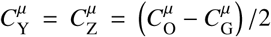. The GCCs of all other proteins were sampled from a probability distribution that respects equation (11) and assumes that proteins with a larger abundance tend to have a larger GCC (see S1 Text, p. 11

## Acknowledgements

We thank Daan Kiviet and Philippe Nghe for sharing the cross-correlation data that was used to generate Fig 4DEF, top panels.

This work was supported by the NWO (Nederlandse Organisatie voor Wetenschappelijk Onderzoek) (Grant 022.005.023).

